# Sertraline modulates hippocampal plasticity and learning via sigma 1 receptors, cellular stress and neurosteroids

**DOI:** 10.1101/2024.01.23.576911

**Authors:** Yukitoshi Izumi, Angela M. Reiersen, Eric J. Lenze, Steven J. Mennerick, Charles F. Zorumski

**Author notes:** **Correspondence:** Charles. F. Zorumski, M.D., Department of Psychiatry, Washington University School of Medicine, 660 South Euclid Avenue, St. Louis MO 63110.

## Abstract

In addition to modulating serotonin transport, selective serotonin reuptake inhibitors (SSRIs) have multiple other effects that may contribute to clinical effects, and some of these latter actions prompt repurposing of SSRIs for non-psychiatric indications. We recently observed that the SSRIs fluvoxamine and fluoxetine prevent the acute adverse effects of pro-inflammatory stimulation on long-term potentiation (LTP) in the CA1 hippocampal region. Sertraline showed markedly different effects, acutely inhibiting LTP at a low micromolar concentration through inverse agonism of sigma 1 receptors (S1Rs). In the present studies, we pursued mechanisms contributing to sertraline modulation of LTP in rat hippocampal slices. We found that sertraline partially inhibits synaptic responses mediated by N-methyl-D-aspartate receptors (NMDARs) via effects on NMDARs that express GluN2B subunits. A selective S1R antagonist (NE-100), but not an S1R agonist (PRE-084) blocked effects on NMDARs, despite the fact that both S1R ligands were previously shown to prevent LTP inhibition. Both NE-100 and PRE-084, however, prevented adverse effects of sertraline on one-trial learning. Because of the important role that S1Rs play in modulating endoplasmic reticulum stress, we examined whether inhibitors of cellular stress alter effects of sertraline. We found that two stress inhibitors, ISRIB and quercetin, prevented LTP inhibition, as did inhibitors of the synthesis of endogenous neurosteroids, which are homeostatic regulators of cellular stress. These studies highlight complex effects of sertraline, S1Rs and neurosteroids on hippocampal function and have relevance for understanding therapeutic and adverse drug actions.

## INTRODUCTION

Selective serotonin reuptake inhibitors (SSRIs) are among the most widely prescribed psychiatric medications with benefits across mood, anxiety and stress-related disorders [1]. These agents are thought to produce beneficial effects primarily via their ability to inhibit serotonin transport, increasing extracellular serotonin levels and prolonging actions following synaptic release. Among the SSRIs, sertraline is favorably viewed as being free of drug-drug interactions and off-target effects. Yet, SSRIs have multiple additional actions, including intracellular effects that are not dependent upon serotonin [2]. These latter effects include synthesis of neurosteroids, inhibition of acid sphingomyelinase, modulation of sigma-1 receptors (S1Rs), alteration of endoplasmic reticulum (ER) stress responses, stimulation of macroautophagy, and anti-inflammatory effects, among others [3–10]. The pleiotropic actions of SSRIs have made these agents attractive candidates for re-purposing as treatments for a variety of non-psychiatric illnesses [1,11], but also raise the possibility of adverse effects from these non-serotonergic actions.

Recent studies support the use of certain SSRIs in treatment of inflammatory conditions both in preclinical models and in specific patient populations [12–14]. Of particular note, the SSRI, fluvoxamine, has shown beneficial effects in preventing deterioration in individuals with newly diagnosed COVID-19 [12,13]. Initial findings with fluvoxamine have been replicated in some [15,16], but not all studies [17]. Importantly, the potential of fluvoxamine and other SSRIs as modulators of inflammation, including neuroinflammation, coupled with the relative safety and low cost of these agents make them attractive candidates for further study.

In recent experiments, we examined the effects of fluvoxamine, fluoxetine and sertraline in a well-characterized model of acute inflammation in the rodent hippocampus. In this model, lipopolysaccharide (LPS), a bacterial endotoxin and strong pro-inflammatory stimulus, does not alter baseline synaptic transmission in the CA1 hippocampal region, but markedly dampens the ability to induce long-term potentiation (LTP), a form of synaptic plasticity thought to underlie learning and memory [18]. Effects of LPS are prevented by pretreatment with fluvoxamine and fluoxetine via mechanisms that involve activation of S1Rs and stimulation of local hippocampal synthesis of 5-alpha reduced neurosteroids in the case of fluvoxamine, and stimulation of neurosteroid production by an S1R-independent mechanism in the case of fluoxetine [19].

In these initial studies, sertraline, a highly potent SSRI that promotes neurosteroid synthesis [4], exhibited markedly different effects than fluvoxamine and fluoxetine. When administered alone at a low micromolar concentration, sertraline acutely inhibited LTP induction by a mechanism that was reversed by either a selective S1R agonist or S1R antagonist [19]. These latter observations are consistent with prior studies indicating that sertraline is an S1R inverse agonist [6,20,21], but leave unanswered how sertraline produces its effects on hippocampal plasticity. In the present study, we examined potential mechanisms underlying effects of sertraline on hippocampal LTP focusing on the roles of S1Rs and neurosteroid production.

## MATERIALS and METHODS

### Hippocampal slice preparation

The Washington University IACUC approved all protocols for animal experiments. Under isoflurane anesthesia, we prepared hippocampal slices from postnatal day (P) 28-33 Harlan Sprague-Dawley male albino rats (Indianapolis IN) using standard methods [22,23]. Juvenile male rats were used for these studies because of the robust and highly reliable synaptic plasticity at this age and to avoid estrus cycle-related effects. Dissections were done in ice-cold artificial cerebrospinal fluid (ACSF) containing (in mM): 124 NaCl, 5 KCl, 2 MgSO_4_, 2 CaCl_2_, 1.25 NaH_2_PO_4_, 22 NaHCO_3_, 10 glucose, gassed with 95% O_2_-5% CO_2_ at 4-6°C. The dorsal two-thirds of the hippocampus was cut into 500 µm slices with a rotary tissue slicer and maintained in ACSF at 30°C for at least 1 hour before experiments.

### Hippocampal slice physiology

Individual slices were transferred to a submersion-recording chamber and perfused with ACSF at 2 ml/min and 30°C. Extracellular recordings were obtained from CA1 *stratum radiatum* to monitor excitatory postsynaptic potentials (EPSPs) using constant current pulses (0.1 ms) to the Schaffer collateral pathway every minute via a bipolar stimulating electrode. Stimulus intensity was half-maximal based on a baseline input-output (IO) curves. We induced LTP using a single 100 Hz by 1 s high frequency stimulation (HFS) and repeated IO curves 60 min following HFS. For display, responses are typically shown at 5 min intervals. Long-term depression (LTD) was induced using 1 Hz low frequency stimulation (LFS) for 15 min (900 pulses) as described previously [24,25].

NMDAR-mediated synaptic responses were recorded in *stratum radiatum* by stimulation of the Schaffer collateral pathway once per minute. For these experiments, we used ACSF containing 0.1 mM Mg^2+^and 2.5 mM Ca^2+^ to facilitate signaling via NMDARs, and 30 μM 6-cyano-7-nitroquinoxaline-2,3-dione (CNQX) to block AMPAR responses [24]. Residual EPSPs are completely inhibited by a saturating concentration of the selective NMDAR-antagonist, D-2-amino-5-phosphonovalerate (APV) [24,25]. We quantified NMDAR responses as the rising slope of the field potential before and at the end of drug perfusion.

### Behavioral Studies

We examined acute effects of sertraline on learning and memory using a one-trial inhibitory avoidance task that is dependent upon hippocampal plasticity [18,22,26]. In this task, rats are placed in a chamber that has lit (1000 lux) and dark (< 10 lux) compartments. Both compartments have floors of stainless steel rods, through which an electrical shock can be administered in the dark chamber. On the first day of training, rats were placed in the lit chamber and allowed to move freely between chambers for 10 min. One hour later, rats were administered either saline, PRE-084, an S1R agonist, or NE-100, an S1R antagonist, at a dose of 10 mg/kg i.p. (dissolved in saline). At hour 2 rats were administered vehicle (DMSO) or sertraline (10 mg/kg i.p. dissolved in 2% DMSO). At hour 3, animals were placed in the lit compartment and allowed to explore the apparatus freely for up to 300 s (5 min). When rats entered the dark chamber, they were given a single foot shock and removed from the apparatus. On the second day of study, rats were again placed in the lit chamber and the time spent in the light was recorded over a 300 s trial. No foot shocks were administered on the second day of study.

### Chemicals

Sertraline (CAS#:79559-97-0), PRE-084 (CAS#:138847-85-5), dutasteride (CAS#:164656-23-9) and LPS were purchased from Millipore Sigma (St. Louis MO), as were salts. NE-100 (CAS#:149409-57-4) was purchased from Tocris Bioscience (Ellisville MO). Finasteride (CAS#:98319-26-7) was from Steraloids (Newport RI).

### Data Collection & Analysis

Electrophysiology results were acquired and analyzed using pClamp (Molecular Devices, Union City CA). Results in the text display mean ± SEM with responses normalized to baseline (100%). Statistical analyses are based on comparison of IO curves obtained at baseline and 60 min following HFS [22–26]. Student’s t-tests were used for most comparisons, except as noted in the text. For studies examining changes in NMDA EPSPs (Figure 1B-C), data were analyzed by repeated measures ANOVA followed by Tukey’s multiple comparison test. Behavioral studies with multiple comparisons were analyzed using the Kruskal-Wallis test followed by Dunn’s multiple comparison test (GraphPad Prism 9.2.0, GraphPad Software, La Jolla CA). Values in the text reflect the number (N) of animals in a condition. SigmaStat (Systat Software, Inc., Richmond City, CA) was used for other comparisons. Synaptic responses shown in figures are from continuous monitoring of responses at low frequency stimulation and may differ from numerical results in the text, which are based on analysis of IO curves.

**Figure 1.**
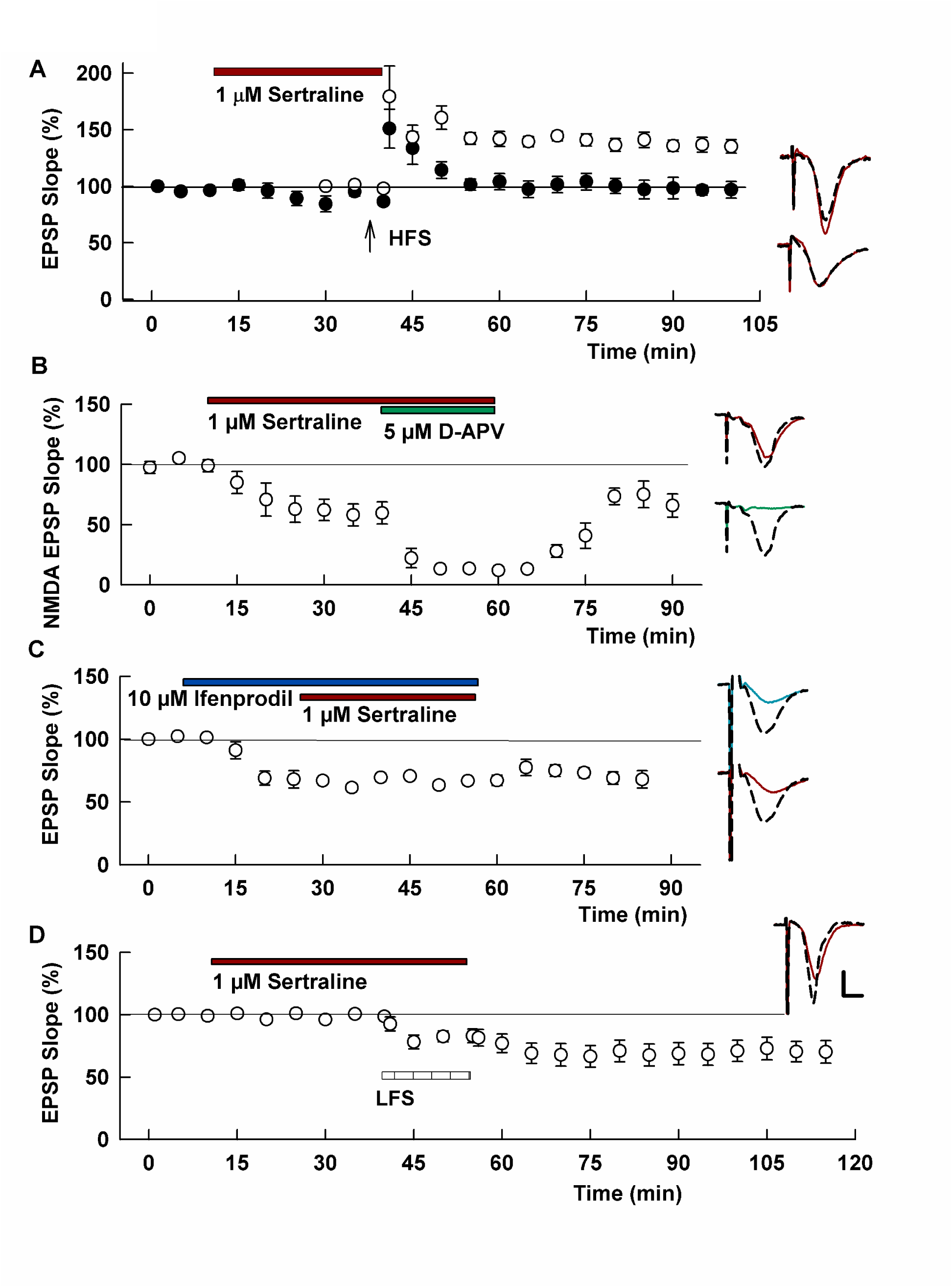
Sertraline inhibits LTP and partially depresses NMDAR responses. A. In control slices, a single 100 Hz x 1 s HFS (arrow) reliably induces LTP in the CA1 region (white circles). A 30 min administration of 1μM sertraline (black bar) completely inhibits LTP induction (black circles). B. Sertraline partially depresses NMDAR-mediated EPSPs. Addition of a low concentration of D-APV (5 μM) produces additive depression of NMDAR responses. C. The GluN2B inhibitor, ifenprodil (10 μM) partially depresses NMDAR responses and addition of sertraline results in no further depression. D. Despite inhibiting GluN2B NMDARs, sertraline failed to block induction of LTD by 1 Hz LFS (hashed bar). Traces to the right of the graph show representative EPSPs under baseline conditions (dashed traces) and 60 min following HFS (A) or LFS (D) (solid traces). In panels B and C, baseline NMDAR EPSPs are dashed and solid traces are in the presence of drugs. Scale bar: 1 mV, 5 ms.

## RESULTS

In the CA1 region of hippocampal slices from juvenile rats, a single 100 Hz x 1 s HFS delivered to the Schaffer collateral pathway reliably induces stable LTP (132.2 ± 1.5%, N=5, Figure 1A). As we showed recently [19], acute administration of 1 μM sertraline for 30 min prior to HFS had no significant effect on baseline synaptic transmission but completely inhibited induction of LTP (100.4 ± 4.5%, N=5; p = 0.0002 vs. control LTP; Figure 1A). In the experiments that follow, we examined potential mechanisms contributing to this negative modulation of hippocampal plasticity.

Because our prior studies found that two other high potency SSRIs [19], fluvoxamine and fluoxetine, had no effect on induction of CA1 LTP in control slices, we focused on non-serotonin mechanisms that could contribute to LTP inhibition. NMDAR activation is critically important for LTP induction under the conditions of our experiments; thus, we initially examined effects of sertraline on isolated NMDAR-mediated EPSPs. NMDAR EPSPs are stable under control conditions with no change in response over more than 40 min of recording (107.6 ± 4.5% of baseline, N=6; Supplemental Figure 1). In the presence of sertraline, NMDAR synaptic responses decreased by over 30% (66.9 ± 9.2% of baseline, N=5, p=0.0022 vs. baseline; Figure 1B). Addition of a low concentration of the competitive NMDAR antagonist, D-2-amino-5-phosphovalerate (APV, 5 μM) that partially inhibits NMDARs [24,25] produced complementary suppression of these responses resulting in over 90% total inhibition (9.7 ± 1.3% of baseline, p = 0.0159 vs. sertraline alone by Tukey’s test, N=5). Following washout of both sertraline and APV for 30 min, responses only partially recovered to 62.0 ± 11.7% of baseline (N=5; p=0.5317 by Tukey’s test), similar to the degree of suppression of NMDAR responses in the presence of sertraline alone. The results suggest a persisting inhibition of NMDAR function by sertraline.

The complementary inhibition of NMDAR responses by sertraline and low APV prompted us to examine whether sertraline preferentially alters a subclass of NMDARs bearing GluN2B subunits, as we have observed previously with ethanol and ketamine [24,27]. We found that a saturating concentration of the GluN2B selective inhibitor ifenprodil (10 μM) blocked NMDAR EPSPs by 30-40% akin to sertraline (63.1 ± 6.9% of baseline responses, N=6, p=0.0003 vs. baseline; Figure 1C). In the presence of ifenprodil, sertraline produced no further inhibition (66.0 ± 7.0% of baseline, N=6; p=0.1706 vs. ifenprodil; Figure 1C). These results indicate that sertraline is a partial but persistent inhibitor of NMDARs at a concentration that blocks LTP completely, acting largely at a GluN2B-containing subclass of receptor. In prior studies, we found that similar degrees and type of NMDAR inhibition alone are insufficient to block LTP induction [24,25,27]. Additionally, we have found that LTD induced by 1 Hz low frequency stimulation (LFS) involves GluN2B NMDARs and is inhibited by ifenprodil [24,25]. In the presence of sertraline, however, LTD could still be induced with some variability among slices (67.8 ± 8.2% of baseline 60 min following LFS, N=9; Figure 1D; p=0.9916 vs. control LFS-induced LTD: 68.0 ± 5.0%, N=5; Supplemental Figure 2).

Sertraline differs from other SSRIs, particularly fluoxetine and fluvoxamine, in functioning as an inverse agonist/antagonist at S1Rs [20,21]. Furthermore, S1Rs modulate NMDARs [28–33] and we previously found that both a selective antagonist and agonist of S1Rs overcomes the ability of sertraline to inhibit LTP [19]. At 1 μM, the selective S1R antagonist, NE-100 [34], had no effect on NMDAR EPSPs alone (102.7 ± 6.5% of baseline, N=5) but completely inhibited the effects of sertraline (97.9 ± 6.9%; p = 0.5484 vs. NE-100 alone by paired t-test, Figure 2A). Addition of 5 μM APV blocked NMDAR responses by about 50% (49.3 ± 6.0% of baseline, N=5; p = 0.0047 vs. NE-100 + sertraline). Following washout of all three drugs, responses recovered to 83.3 ± 10.5% (N=5; p=0.2027 vs. sertraline + NE-100) of baseline, not differing from sertraline plus NE-100.

**Figure 2.**
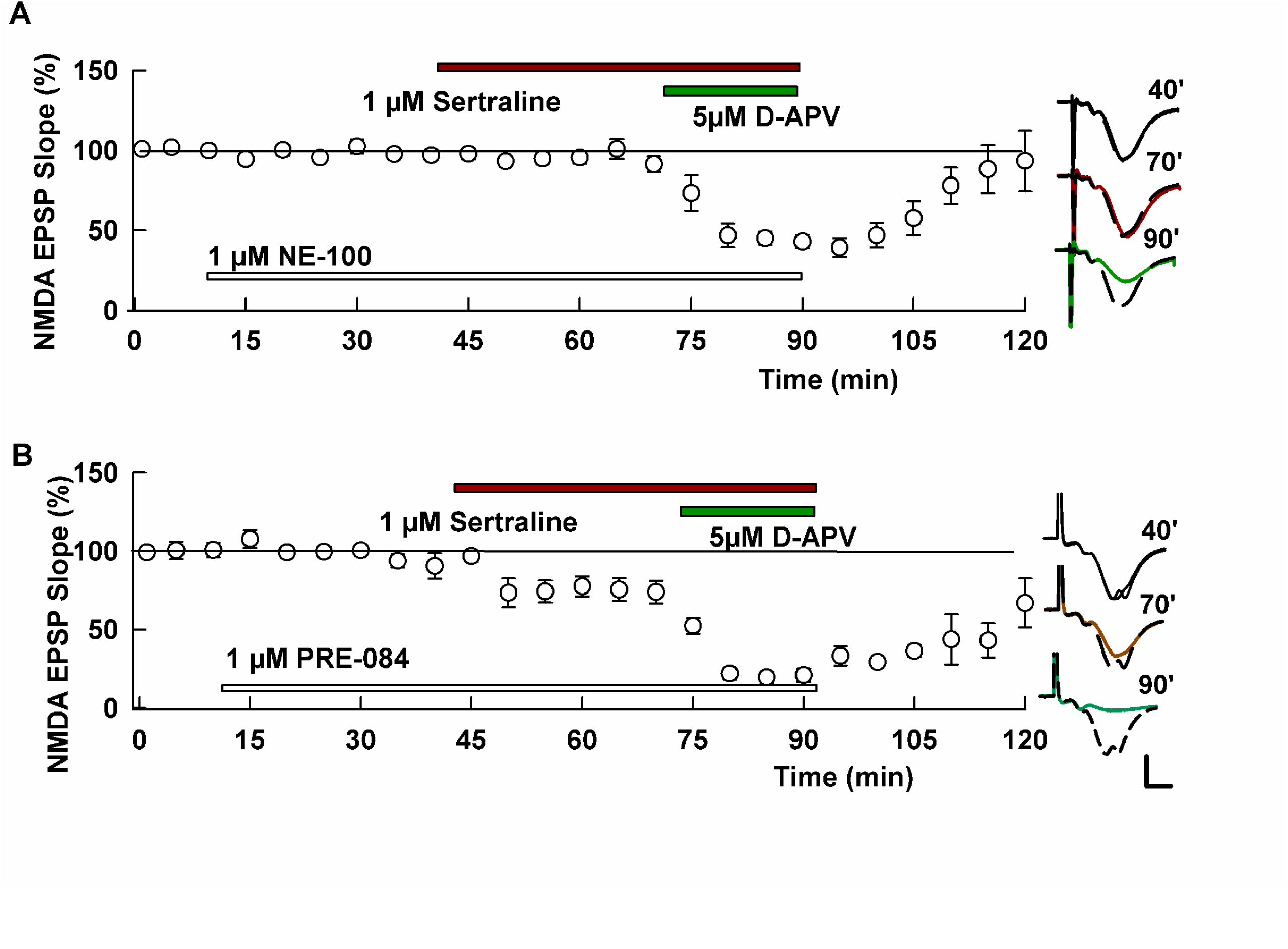
Effects of sertraline on NMDAR responses involve S1Rs. A. At 1 μM, the S1R antagonist NE-100 (white bar) had no effect on NMDAR EPSPs but blocked the effects of sertraline (black bar). A low concentration of APV (as in Figure 1, gray bar) resulted in partial suppression of these responses. B. The S1R agonist, PRE-084 (1 μM), a concentration that overcame the effects of sertraline on LTP in our prior study (Izumi et al., 2023) had no effect on NMDAR responses alone and failed to inhibit the effects of sertraline. As in Figure 1, low APV produced additive depression of responses. Traces show representative NMDAR EPSPs at the times indicated as in Figure 1. Calibration bar: 1 mV, 5 ms.

In contrast to the S1R antagonist, 1 μM PRE-084, a selective S1R agonist [35] that we showed previously overcame the effects of sertraline on LTP [19], had no effect on NMDAR responses when administered alone (97.6 ± 2.7% of baseline, N=5). PRE-084 also did not prevent inhibition of NMDAR responses by sertraline (57.8 ± 10.3% of baseline, N=5, p=0.0123 vs. PRE-084 alone by paired t-test; Figure 2B). Addition of APV depressed these responses to 13.9 ± 3.8% of baseline (N=5; p=0.0047 vs. sertraline + PRE-084). Recovery of responses following washout of these three agents was again poor and consistent with what was observed with sertraline alone (32.9 ± 11.9%, N=5 of baseline; p=0.2093 vs. sertraline + PRE-084). These results indicate that sertraline is a partial inhibitor of NMDARs at a concentration that blocks LTP completely, and that an S1R antagonist but not S1R agonist prevents this inhibition.

The ability of a partial inhibitor of NMDARs to block CA1 LTP is reminiscent of what we have observed previously with other non-competitive NMDAR antagonists including ethanol [23,36,37] and ketamine [27]. In those cases, the NMDAR inhibitor produces a form of metaplastic LTP block that persists after drug washout and is inhibited by complete NMDAR inhibition with a high concentration of APV during the period of drug exposure, with washout of both agents for at least 30 min prior to HFS [36]. To test whether similar metaplasticity occurs with sertraline, we administered the SSRI for 30 min followed by 30 min drug washout prior to delivery of an LTP-inducing tetanus. Under these conditions, sertraline alone produced persisting LTP inhibition (96.5 ± 4.0% of baseline 60 min following HFS, N=5, p = 0.0001 vs. control LTP; Figure 3A), but, unlike ethanol and ketamine, this LTP inhibition was not prevented by a saturating concentration of APV administered during sertraline administration (103.2 ± 4.1% of baseline, N=5, p = 0.2825 vs. sertraline alone; Figure 3B).

**Figure 3.**
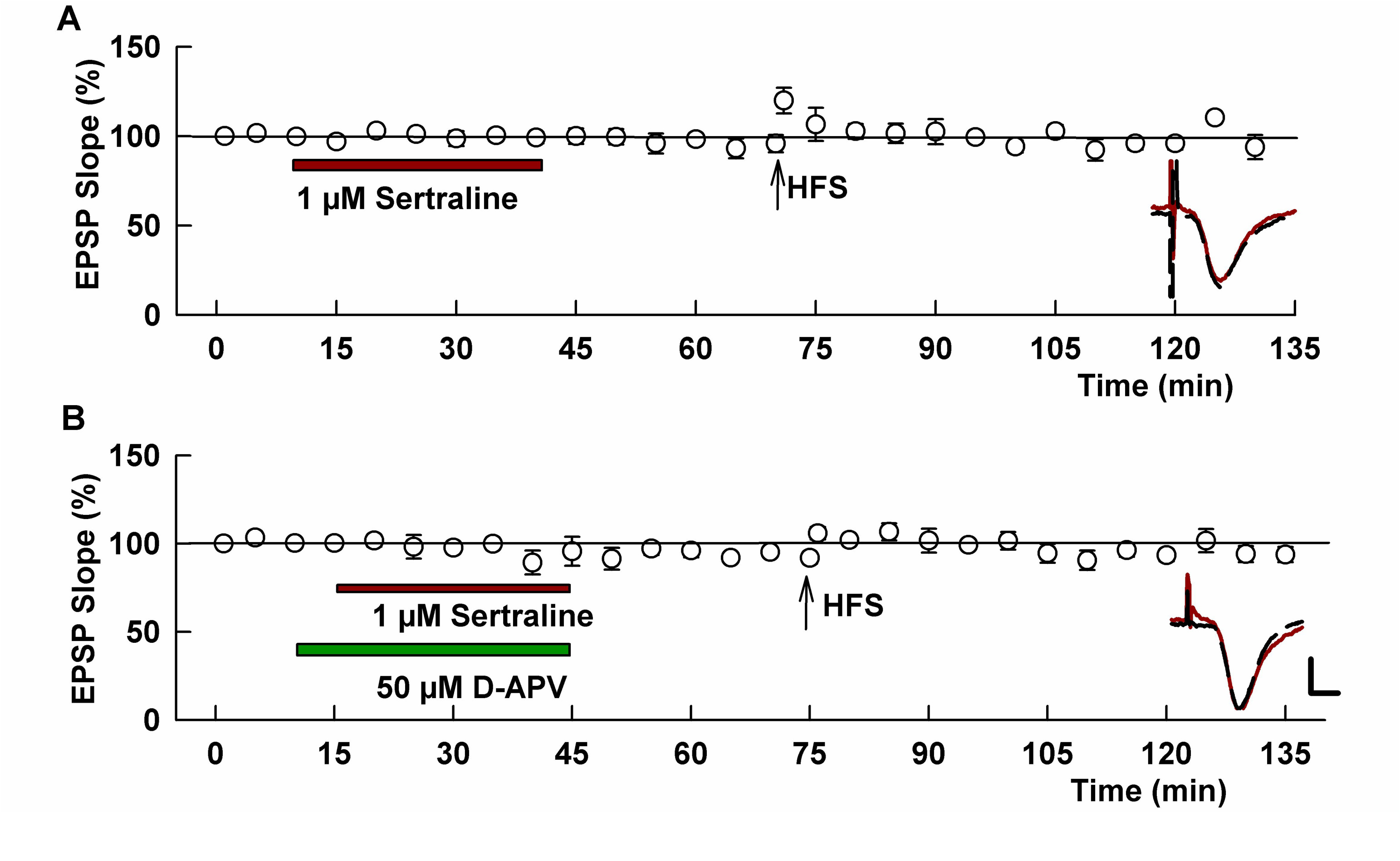
Sertraline administration results in persistent LTP inhibition that is not altered by co-administration of saturating APV. A. Administration of sertraline for 30 min (black bar) followed by 30 min washout prior to HFS (arrow) resulted in complete suppression of LTP induction. B. In contrast to what we have observed with agents that produce NMDAR-dependent metaplastic LTP inhibition [36], administration of sertraline in the presence of saturating APV (50 μM, gray bar) followed by washout of both agents 30 min prior to APV did not prevent LTP inhibition. Traces show EPSPs as in Figure 1A. Calibration: 1 mV, 5 ms.

S1Rs are molecular chaperones synthesized in endoplasmic reticulum (ER) and expressed in mitochondrial associated membranes (MAMs) where they help to regulate ER and mitochondrial function [38–40]. Furthermore, ER stress and the cellular integrated stress response (ISR) are known regulators of hippocampal synaptic plasticity [41–43], and sertraline is reported to promote ER stress [44,45]. Thus, we examined whether agents that inhibit ER stress and the ISR modulate the effects of sertraline on CA1 LTP. We found that the ISR inhibitor, ISRIB (1 μM) [42], prevented the effects of sertraline on LTP when administered prior to and during sertraline exposure (129.3 ± 1.9% of baseline, N=5; p = 0.0004 vs. sertraline alone; Figure 4A). When administered alone, ISRIB had no effect on LTP induction under control conditions [41]. We also examined the effects of quercetin (50 μM), an agent that has antioxidant properties and inhibits ER stress [46–48]. Quercetin pre-treatment, like ISRIB, prevented the effects of sertraline on LTP (133.4 ± 4.5% of baseline, N=5; p=0.0008 vs. sertraline; Figure 4B), but had no effect on LTP induction under control conditions (132.8 ± 5.7%, N=5; p=0.9214 vs. control LTP; Supplemental Figure 3).

**Figure 4.**
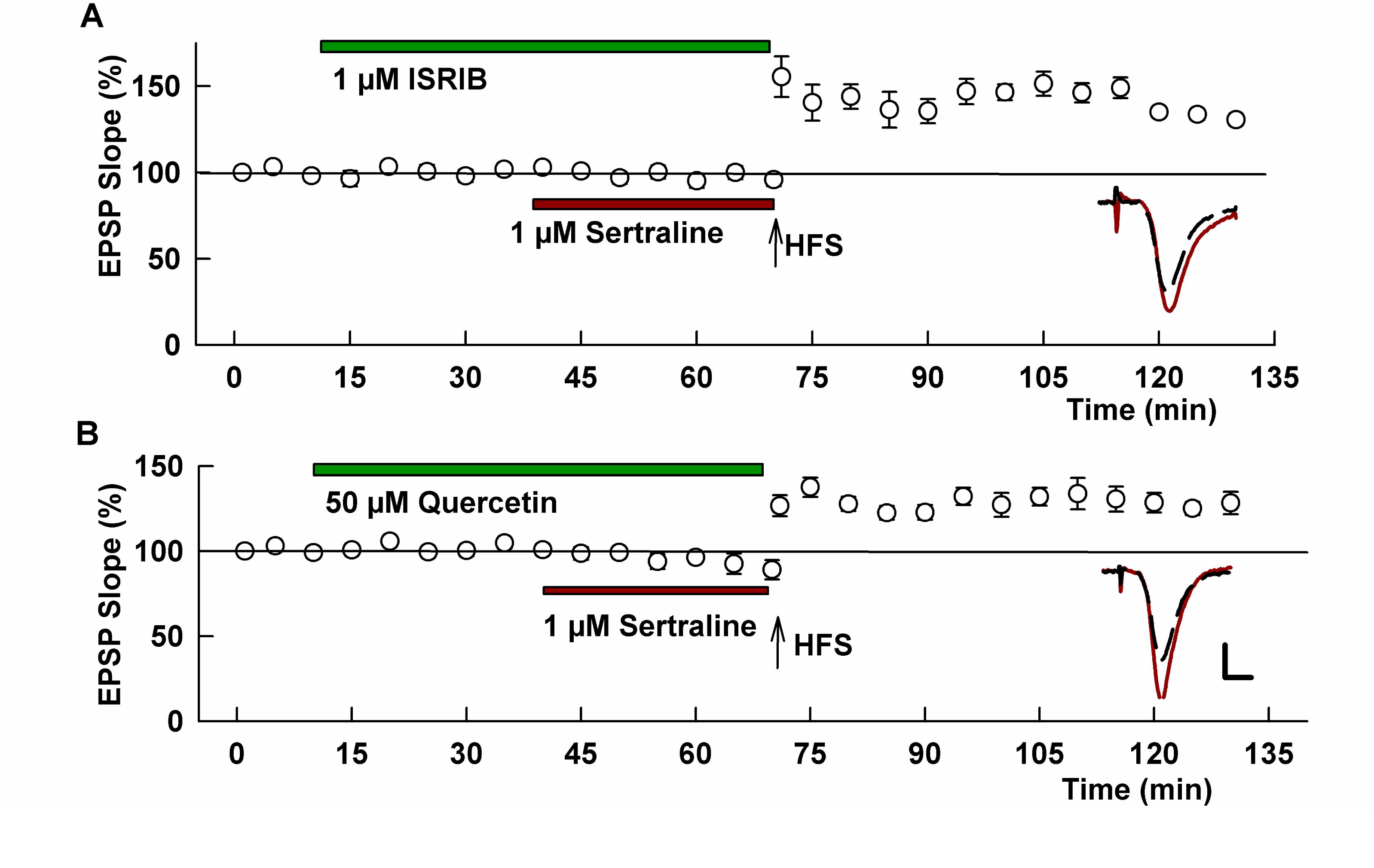
Inhibitors of cellular stress prevent the effects of sertraline on LTP. A. Administration of 1 μM ISRIB (green bar), an agent that inhibits the integrated stress response, overcame the acute effects of sertraline (red bar) on LTP. B. Quercetin (50 μM, green bar), which inhibits ER stress, also prevented the effects of sertraline (red bar) on LTP when perfused before and during sertraline. Traces show representative EPSPs; dashed traces are baseline and red traces are 60 min after HFS. Scale bar: 1 mV, 5 ms.

The role of cellular stress responses in the effects of sertraline raise questions about endogenous homeostatic mechanisms that could contribute to LTP inhibition. In the face of ER stress, cholesterol is mobilized from ER and results in synthesis of neurosteroids as a mechanism to regulate stress responses and preserve neuronal integrity [49]. Additionally, sertraline has direct effects on enzymes involved in the synthesis of GABAergic 5α-reduced neurosteroids most prominently allopregnanolone (AlloP) [4], and we previously found that endogenous 5α-steroids negatively modulate CA1 plasticity under stressful conditions [36,37]. To examine a potential role of 5α-reduced neurosteroids in the effects of sertraline, we used two well-characterized and selective inhibitors of 5-alpha reductases (5AR), finasteride and dutasteride [50–52]. In prior studies, we found that neither of these 5AR inhibitors altered CA1 LTP under baseline conditions [19,22,23]. In slices treated with 1 μM finasteride, a 5AR inhibitor that preferentially affects Type II 5AR [53,54], sertraline-induced LTP inhibition was not altered (85.3 ± 7.2%, N=5; p=0.1132 vs. sertraline alone; Figure 5A). However, 1 μM dutasteride, a broader spectrum and more persistent 5AR inhibitor [50–53,55,56], completely prevented LTP inhibition by sertraline (150.0 ± 15.4% of baseline, N=5; p=0.0149 vs. sertraline alone; Figure 5B). Results with dutasteride prompted us to examine a higher concentration of finasteride because finasteride’s effects on Type I 5AR are competitive and more readily reversible [53]. At 10 μM, finasteride, like dutasteride, completely prevented the effects of sertraline on LTP (145.2 ± 9.5%, N=5, p=0.0028 vs. sertraline alone; Figure 5A), although we did observe some depression of baseline responses in the presence of finasteride plus sertraline.

**Figure 5.**
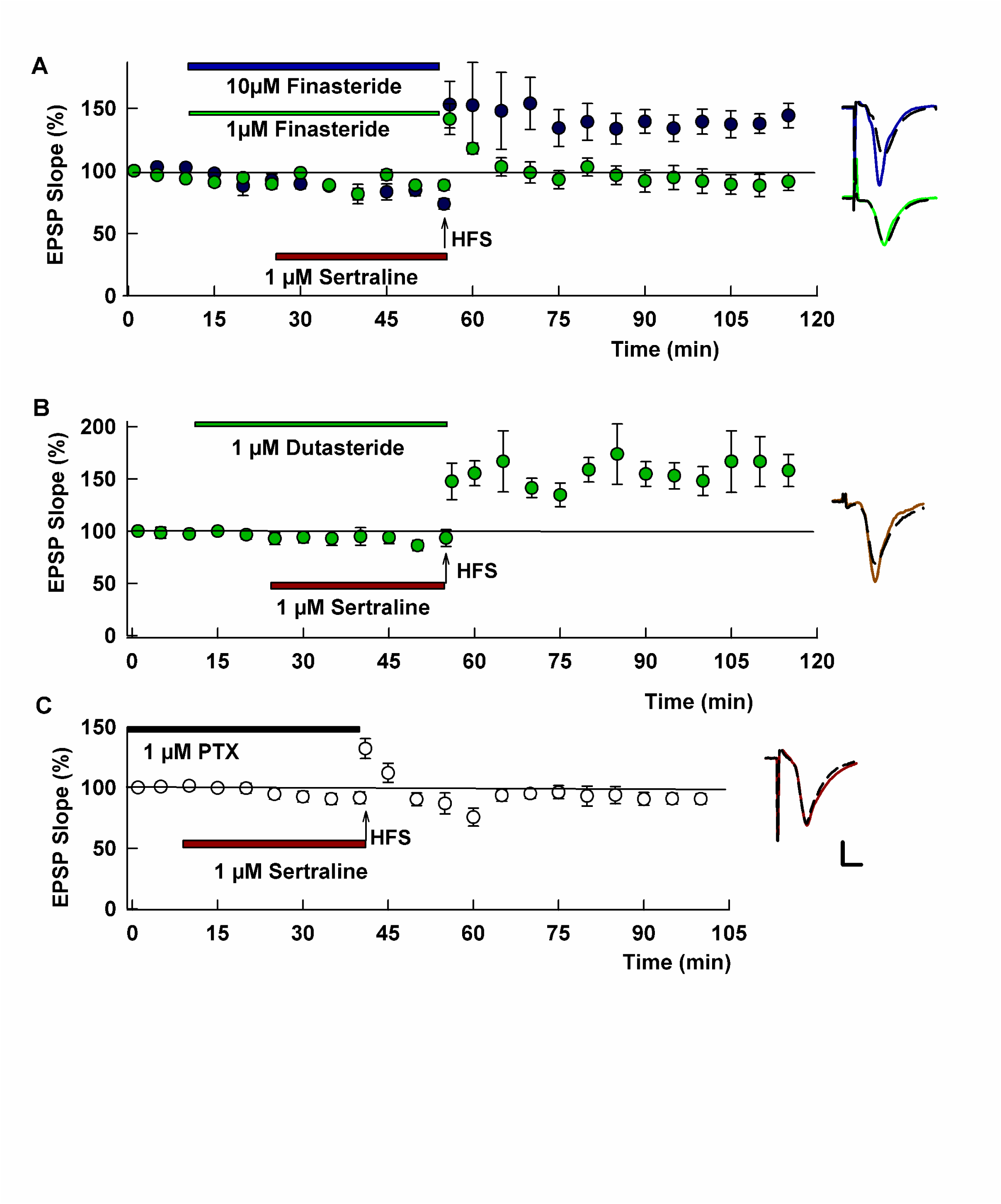
Inhibitors neurosteroid synthesis prevent the effects of sertraline on LTP. A. The 5-alpha reductase (5AR) inhibitor finasteride prevented the effects of sertraline on LTP when administered at 10 μM (dark green) but not 1 μM (light green). B. The more potent broader spectrum 5AR antagonist, dutasteride (1 μM), completely prevented the effects of sertraline when perfused before and during sertraline. C. At 1 μM, the GABA_A_R antagonist picrotoxin (PTXN) failed to alter LTP inhibition by sertraline. Traces show EPSPs. Calibration: 1 mV, 5 ms.

Certain 5-alpha reduced neurosteroids, including AlloP, positively modulate GABA-A receptors and augmented GABAergic inhibition contributes to metaplastic LTP inhibition [36]. Thus, we also examined whether picrotoxin (PTXN), a non-competitive GABA-A receptor antagonist, alters sertraline’s effects on LTP. We found, however, that 1 μM PTXN, which prevented endogenous neurosteroid effects in our prior *ex vivo* studies [23,56–58] had no effect on LTP inhibition (98.7 ± 4.1%, N=5; p=0.7879 vs. sertraline alone; Figure 4C). A higher concentration of PTXN (3 μM) also had no effect (91.9 ± 7.8%, N=5), and at 10 μM PTXN slices displayed epileptiform changes (not shown). Taken together, results in Figures 4 and 5 indicate that cellular stress responses and activation of homeostatic protective responses via 5α-reduced neurosteroids contribute to the effects of sertraline on LTP, but the effects of neurosteroids are unlikely mediated by altered GABAergic inhibition.

Given the effects of sertraline on LTP induction and the role of LTP in memory processing, we also examined acute effects of the drug on a form of inhibitory avoidance learning that has been associated with CA1 hippocampal LTP [18,22,25]. In this task, rodents learn to avoid the dark chamber of a two-chamber device following a single foot shock in the dark chamber during conditioning. Control rats administered vehicle readily learn the task and remain in the lit chamber for the full 5 min testing period one day after conditioning (Figure 6). In contrast, rats treated with 10 mg/kg ip sertraline 1 hour prior to conditioning showed variable but significant deficits in learning, remaining in the light for 151.0 ± 43.1 sec (p=0.0064 by Kruskal-Wallis test followed by Dunn’s test for multiple comparisons; Figure 6). Pretreatment of animals 1 h prior to sertraline before conditioning with 10 mg/kg ip of either PRE-084 or NE-100 completely prevented the adverse effects of sertraline (p=0.0064; Figure 6), consistent with their ability to prevent effects of the SSRI on LTP induction [19].

**Figure 6.**
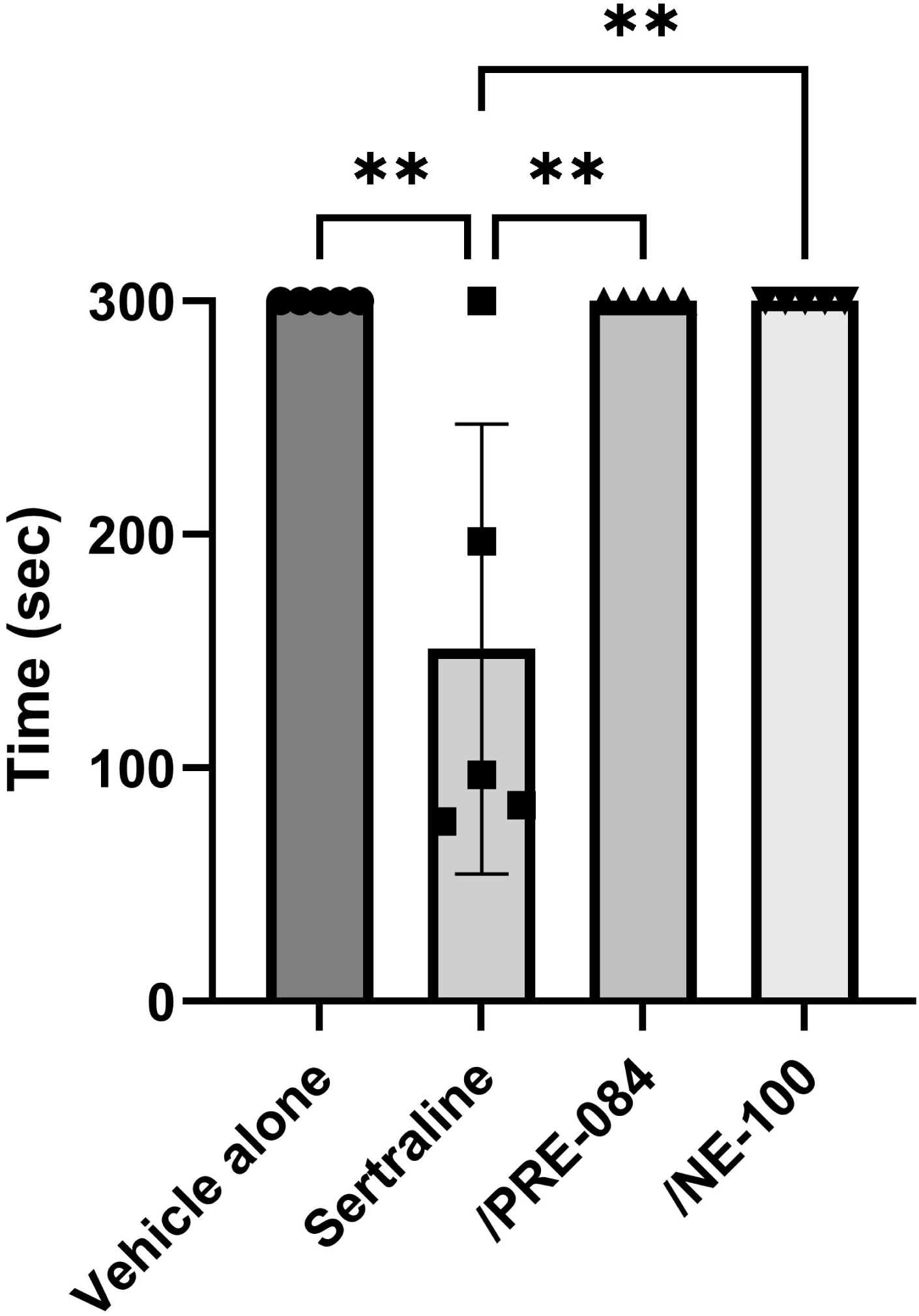
Sertraline acutely dampens one-trial inhibitory avoidance learning in an S1R-dependent fashion. The graph shows that control rats treated with vehicle (DMSO) alone readily learn the task and remain in the lit compartment for the full 300 s trial when tested 24 h following conditioning. Sertraline (10 mg/kg ip) administered an hour prior to conditioning produced defects in learning that were completely reversed by either an S1R agonist (PRE-084,10 mg/kg ip) or S1R antagonist (NE-100, 10 mg/kg ip) administered 1 h prior to sertraline. ** p = 0.0064 by Dunn’s multiple comparison test following Kruskal-Wallis test.

## DISCUSSION

SSRIs are important neurotherapeutics with diverse effects on brain function. In addition to antidepressant and anxiolytic actions likely mediated by serotonin, these agents have multiple other cellular targets that contribute to modulation of inflammation, cellular stress, autophagy and neuroplasticity, including effects on S1Rs and neurosteroidogenesis [1,9]. SSRIs are lipophilic weak bases that readily access intracellular compartments and have direct interactions with lipid membranes and intracellular proteins [2,59–61].

We recently examined acute effects of three SSRIs, fluvoxamine, fluoxetine and sertraline, on hippocampal function in a well-characterized model of neuroinflammation using the bacterial endotoxin, LPS, an agent that strongly activates microglia and pro-inflammatory signaling. While LPS alone had little acute effect on basal transmission in the CA1 region, it markedly impaired LTP induction, a form of plasticity thought to underlie learning and memory, and impaired hippocampal-dependent learning *in vivo* [18]. Fluvoxamine and fluoxetine acutely prevented these effects of LPS via activation of S1Rs in the case of fluvoxamine and local hippocampal synthesis of 5-alpha reduced neurosteroids by both SSRIs [19]. Sertraline was an outlier in these studies and inhibited CA1 LTP acutely via a mechanism involving inverse agonism at S1Rs. Effects of the three SSRIs were observed at low micromolar concentrations that are achieved in brain with clinical dosing [62–64].

In our present studies, we sought to determine how sertraline negatively modulates LTP, focusing initially on NMDAR-mediated responses because these receptors play a critical role in LTP induction [36]. We found that sertraline dampens NMDAR EPSPs by 30-40% and this inhibition is additive with a low concentration of the competitive NMDAR antagonist, APV, but occluded by saturating ifenprodil, a selective inhibitor of GluN2B expressing NMDARs. It is unclear, however, whether effects on NMDARs play a critical role in LTP inhibition by sertraline because we previously found that ifenprodil does not alter LTP induction under the conditions of our experiments [24,25]. Additionally, sertraline, unlike ifenprodil, does not block NMDAR-dependent homosynaptic LTD.

Because sertraline binds S1Rs and SIR ligands modulate the function and expression of NMDARs [28–31,65], we examined the role of S1Rs in NMDAR inhibition using a selective S1R antagonist (NE-100) and agonist (PRE-084). Prior studies found that sertraline functions as an S1R inverse agonist in assays of neurite extension [6,20,21] and, unlike fluvoxamine, fails to overcome phencyclidine-induced cognitive defects *in vivo* [66]. Consistent with these findings, we previously observed that adverse effects of sertraline on LTP are prevented by both NE-100 and PRE-084 [19]. In our present studies, we found that NE-100 prevented inhibition of NMDAR responses by sertraline without altering those responses itself, consistent with a prior report describing the effects of a different S1R inverse agonist [28]. The S1R agonist, PRE-084, however, failed to alter NMDAR inhibition by sertraline, at a concentration that completely prevented the effects of sertraline on LTP in our prior study [19]. Interestingly, both PRE-084 and NE-100 prevented adverse effects of sertraline on one-trial learning in our present studies. Taken together, these results support the idea that sertraline is a functional inhibitor of NMDARs via inverse agonism at S1Rs, but raise important questions about the role of NMDAR antagonism in its LTP inhibition.

We have previously found that certain non-competitive NMDAR antagonists, including ethanol and ketamine [24,27], inhibit CA1 LTP at concentrations that only partially inhibit NMDARs via GluN2B-expressing receptors, similar to sertraline. With both ethanol and ketamine, LTP inhibition is prolonged and persists beyond the period of drug application. This persistent LTP inhibition involves metaplastic changes resulting from paradoxical activation of NMDARs that are unblocked by the drugs during acute administration, rather than direct NMDAR inhibition. Co-administration of saturating APV with ethanol or ketamine followed by washout of the drugs prior to tetanic stimulation results in robust LTP induction [23, 24,27]. Sertraline, however, differs from ethanol and ketamine, and saturating APV failed to prevent its persistent LTP inhibition. Recent studies indicate that SSRIs are cleared within minutes from neurons following drug removal, so persistent LTP block by sertraline is unlikely to involve prolonged presence of the drug within cells or membranes [61], but could involve other lasting drug effects beyond those identified here.

We have also observed persistent metaplastic LTP inhibition under conditions that activate neuronal stress responses including brief bouts of hypoxia and low glucose among others [36]. In these conditions, the initial stressor triggers intracellular signaling that involves activation of serine phosphatases, nitric oxide synthase, p38 MAP kinase [36,67] and the ISR [41]. Inhibition of cellular stress signaling prevents the effects of these neuronal stressors on LTP induction. In the present experiments, we used two approaches to dampen neuronal stress, ISRIB, a specific inhibitor of the ISR [42], and quercetin, a flavenoid that inhibits ER stress [46–48]. Both agents prevented the adverse effects of sertraline on LTP. These observations are consistent with prior studies indicating that sertraline can induce ER stress [44,45] and that S1Rs play a key role in modulating ER stress [38,68]. Effects on cellular stress may also account for the beneficial effects of the S1R agonist, PRE-084, on LTP but not NMDARs in the presence of sertraline [38,41].

Results presented here indicate that sertraline inhibits LTP induction by inverse agonism at S1Rs resulting in partial NMDAR depression and activation of cellular stress responses. In turn, cellular stress stimulates neuroprotective homeostatic responses to preserve neuronal integrity. Among these homeostatic mechanisms is mobilization of cholesterol from ER to mitochondria, with side chain cleavage to pregnenolone, the first step in synthesis of 5-alpha reduced neurosteroids including AlloP [49]. AlloP is a potent neuromodulator that enhances inhibition via potentiation of GABA-A receptors and has anti-inflammatory [69,70] and neuroprotective actions [56,57,71]. In prior studies, we found that 5-alpha reduced neurosteroids play a key role in LTP inhibition under conditions of neuronal stress via augmented GABA-mediated inhibition [36]. In our present experiments, we found that adverse effects of sertraline on LTP are prevented by 5-alpha reductase (5AR) inhibitors, but appear to be independent of augmented GABAergic inhibition based on effects of PTXN. While stimulation of S1Rs promotes synthesis of neurosteroids [72], the inverse agonism of sertraline would dampen this effect. However, sertraline also acts directly on 3α-hydroxysteroid dehydrogenase (HSD), a key enzyme in neurosteroidogenesis, promoting the synthesis of AlloP and inhibiting its degradation, with a net result of increasing neurosteroid levels [4, but see 73]. Thus, dual effects of sertraline on neurosteroid production (direct effects on HSD and activation of cell stress responses) may account for the need for a higher concentration of finasteride to reverse LTP inhibition and the lack of effect of low concentrations of PTXN [51–56].

Among the three SSRIs we have studied to date, only sertraline acutely inhibits LTP induction and it does so by unique mechanisms involving S1Rs, NMDARs, cellular stress and neurosteroids. While acute adverse effects on CA1 LTP could cause cognitive dysfunction [74,75], partial NMDAR inhibition and neurosteroid generation may also contribute to therapeutic effects of the drug. Other NMDAR antagonists such as ketamine and nitrous oxide have rapid antidepressant actions at concentrations that only partially inhibit NMDARs [76–78], so effects of sertraline on NMDARs could play a role in therapeutic efficacy. Additionally, AlloP (as brexanolone) is approved for treatment of postpartum depression, and other GABAergic neuroactive steroids are in clinical development for major depression [79]. Sertraline’s cognitive effects in humans have been studied with neuropsychological testing in small clinical trials in depression [80,81]. These studies have not found a negative effect on learning or other neuropsychological functions. In contrast, an observational study of electronic health records found that sertraline use was associated with accelerated cognitive decline among persons with dementia [82]. Thus, the clinical effects of sertraline on cognitive functions, either acutely or chronically, remain insufficiently known.

A limitation of our studies is that we examined acute effects of sertraline in slices from juvenile rodents and effects of NMDAR antagonists on hippocampal transmission differ between juvenile and adult animals [76–78]. Furthermore, chronic dosing with SSRIs has effects that are not readily apparent with acute dosing, including psychotropic effects and augmentation of specific brain synapses [83,84]. In future studies it will be important to understand effects of SSRIs under different experimental and clinical conditions to determine how the pleiotropic effects of these agents contribute to clinical outcomes.

## ACKNOWLEDGEMENTS

Supported by MH101874 (SJM, CFZ), MH123748 (SJM), MH122379 (CFZ, SJM), the Taylor Family Institute for Innovative Psychiatric Research and the Bantly Foundation. The authors thank Kazuko Izumi and Ann Benz for technical assistance, and members of the Taylor Family Institute for helpful comments and advice.

## AUTHOR CONTRIBUTIONS

YI, SJM, AMR, EJL and CFZ conceived the experiments. YI and CFZ designed research and analyzed data. YI performed experiments. CFZ wrote the paper and all authors edited and revised the paper.

## SUPPLEMENTAL FIGURES

**Supplemental Figure 1.** NMDAR-mediated EPSPs triggered by Schaffer collateral pathway stimulation are stable over 40 min of recording. Traces show baseline EPSPs (dashed trace) and responses after 40 min (red trace). Calibration: 1 mV, 5 ms.

**Supplemental Figure 2.** Repeated low-frequency stimulation (LFS, 1 Hz x 900 pulses) induces robust LTD in control slices that is similar in magnitude to LTD induced by LFS in the presence of sertraline. Traces show representative EPSPs as in other figures.

**Supplemental Figure 3.** Quercetin alone does not alter LTP. The graph shows the time course of EPSPs and induction of robust LTP by HFS (arrow). Quercetin (50 μM) was administered during the period denoted by the black bar. Traces show representative EPSPs as in other figures.

