## Supplementary figures and images for "Sertraline modulates hippocampal plasticity and learning via sigma 1 receptors, cellular stress and neurosteroids"

### Supplemental Figure 1

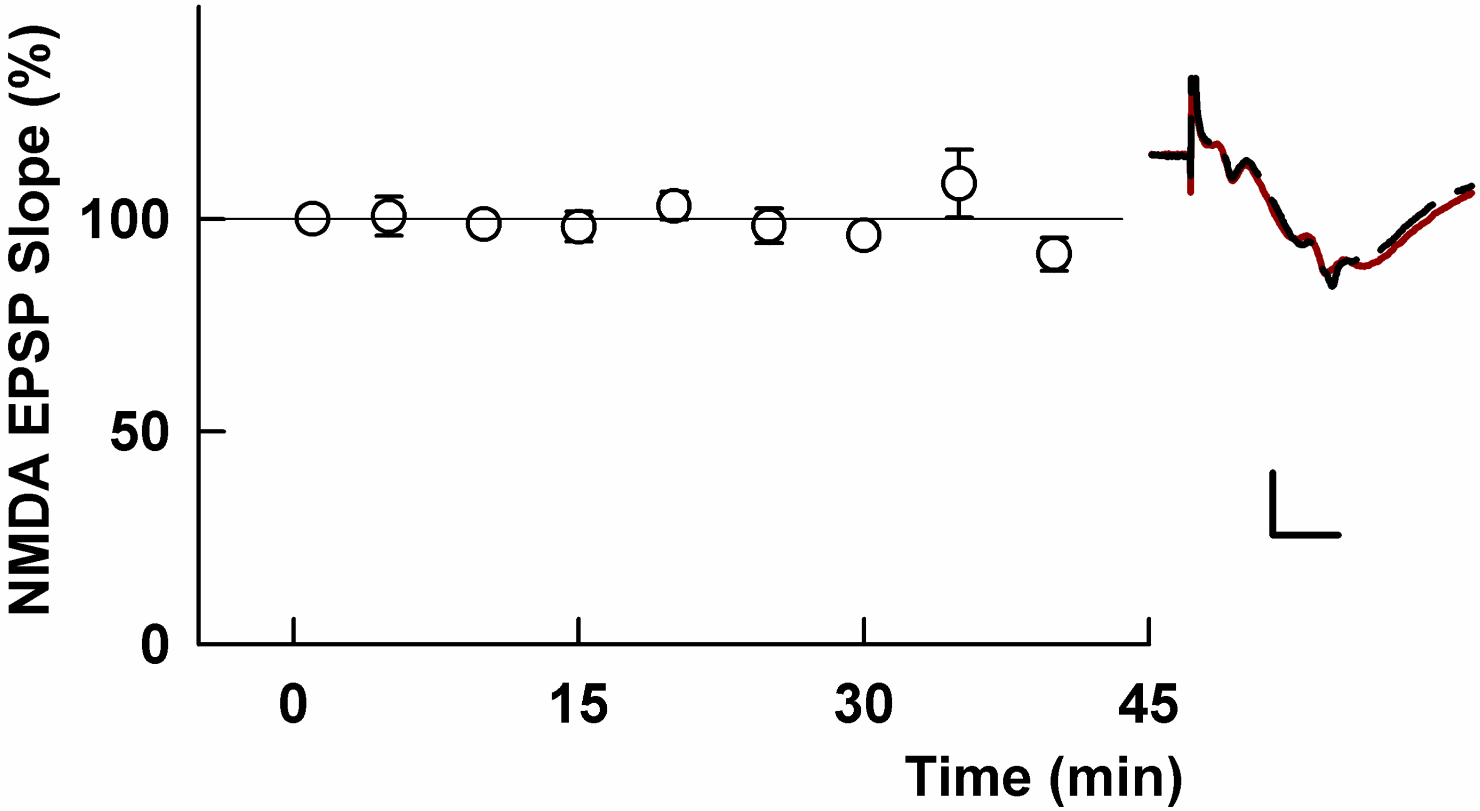

### Supplemental Figure 2

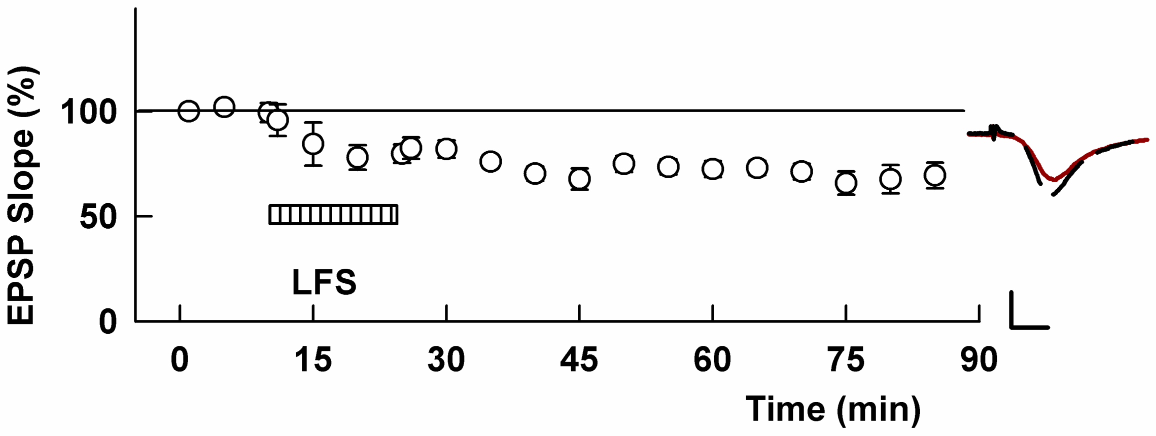

### Supplemental Figure 3

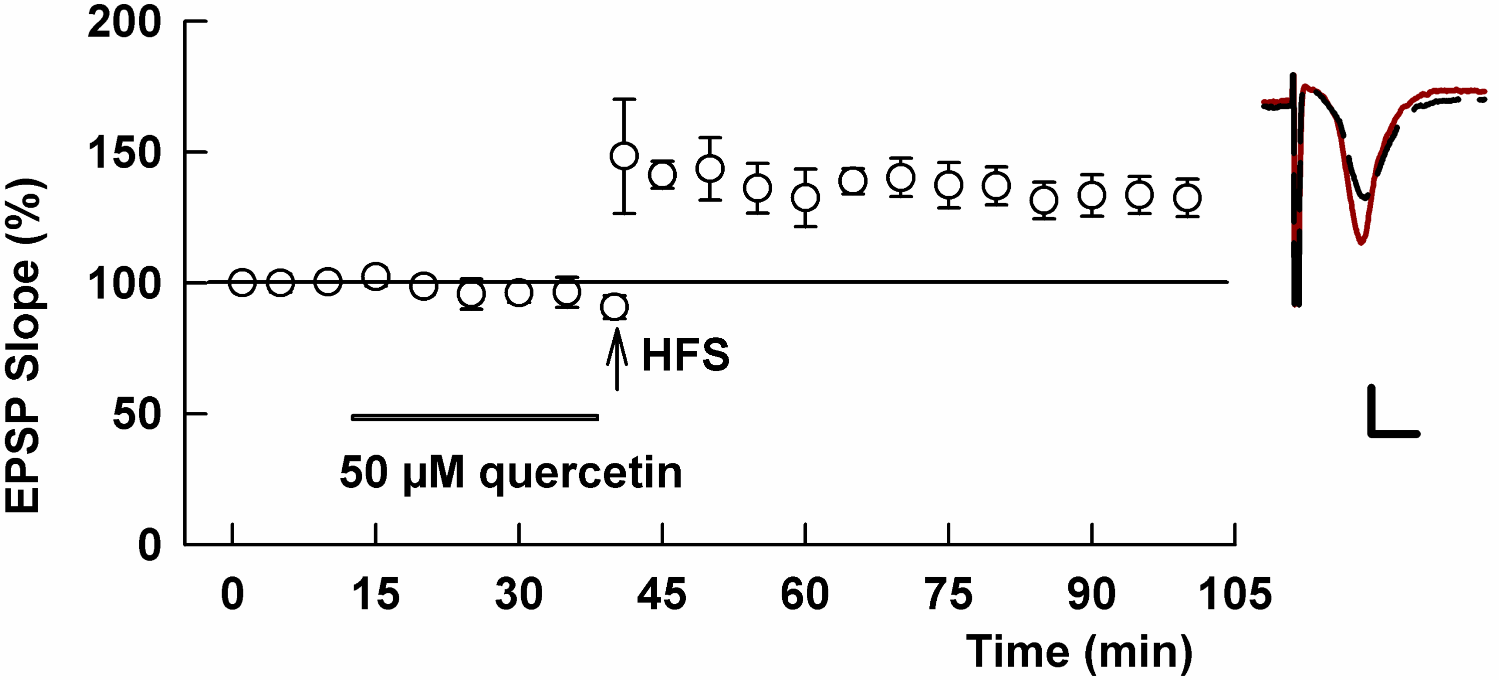
